# Rejuvenating silicon probes for acute electrophysiology

**DOI:** 10.1101/2024.02.20.581222

**Authors:** Alden M. Shoup, Natasha Porwal, Mohammad Amin Fakharian, Paul Hage, Simon P. Orozco, Reza Shadmehr

## Abstract

Electrophysiological recording with a new probe often yields better signal quality than with a used probe. Why does the signal quality degrade after only a few experiments? Here, we considered silicon probes in which the contacts are densely packed, and each is coated with a conductive polymer that increases its surface area. We tested 12 Cambridge Neurotech silicon probes during 61 recording sessions from the brain of 3 marmosets. Out of the box, each probe arrived with an electrodeposited polymer coating on 64 gold contacts, and an impedance of around 50k Ohms. With repeated use, the impedance increased and there was a corresponding decrease in the number of well-isolated neurons. Imaging of the probes suggested that the reduction in signal quality was due to a gradual loss of the polymer coating. To rejuvenate the probes, we first stripped the contacts, completely removing their polymer coating, and then recoated them in a solution of 10 mM EDOT monomer with 32 uM PSS using a current density of about 3mA/cm^2^ for 30 seconds. This recoating process not only returned probe impedance to around 50k Ohms, it also yielded significantly improved signal quality during neurophysiological recordings. Thus, insertion into the brain promoted loss of the polymer that coated the contacts of the silicon probes. This led to degradation of signal quality, but recoating rejuvenated the probes.

## Introduction

Our current ability to simultaneously record extracellular activities of neurons in the brain is due to the development of high-density, multi-contact silicon probes. The large number of contacts makes it possible to record extracellular voltages from not only multiple neurons, but also from multiple regions of a single neuron, thus providing a pseudo image of the geometry of each neuron as well as the spike interactions between neurons.

However, the high density spatial configuration dictates a small contact size (hundreds of micrometers square area) (Ludwig et al., 2006), which results in high impedance. In theory, high impedance can make it more difficult to record the microvolt-range activities of neurons (Loeb et al., 1995; Ludwig et al., 2006; Baranauskas et al., 2011), though the relationship between impedance and recording quality is still unclear (Alba et al., 2015; Neto et al., 2018; Jones et al., 2020). Some probe manufacturers have addressed the impedance issue by using electrodeposition to adhere a conductive polymer on the surface of each contact, increasing the surface area and reducing the impedance (Cui et al., 2001; Ferguson et al., 2009; Niederhoffer et al., 2023). Indeed, contacts with a polymer coating may produce better electrophysiological recordings, but this approach introduces another problem: in acute recordings the probe must be inserted and then retracted, which can remove the polymer coating, thus producing a probe with contacts that have high impedance. The net effect is that a probe with a polymer coating is often useful for only a handful of recordings before it is discarded. Given the high cost of these probes, is there a way to rejuvenate them?

Here, we used 64-contact silicon probes (Cambridge Neurotech) and made impedance and electrophysiological measurements during repeated *in vivo* recordings from the marmoset brain. We found that with repeated use, the probe’s impedance increased, the spike magnitudes decreased, and the number of contacts which failed to record any signals increased. Imaging of the probes suggested that insertion and retraction promoted the removal of the polymer coating. This raised the idea that recoating the contacts via electrodeposition might not only reduce the probe’s impedance, but also restore its signal quality.

To test this idea, we first used an enzyme cleaner to strip the coating from all contacts, and then recoated the contacts using electrodeposition. The results were rejuvenated probes that retained their low impedance and produced high quality recordings that rivaled new probes.

## Results

### PEDOT coating degraded with each recording session, increasing impedance and decreasing signal quality

We used 64-contact silicon probes (M1 checkerboard and M2 linear Cambridge Neurotech) to acquire neurophysiological data from three marmosets. Each session began with an impedance measurement of the probe in saline (see Methods). Next, we inserted the probe through a craniotomy into the brain, traversing the artificial dura (Duragel, Cambridge Neurotech), the intact dura, the visual cortex, and the tentorium, to finally arrive in lobule VI and VII of the cerebellar vermis or the fastigial nucleus (Sedaghat-Nejad et al., 2019). The probe was inserted using a piezoelectric, high precision microdrive (0.5 μm resolution) with an integrated absolute encoder (M3-LA-3.4– 15 Linear smart stage, New Scale Technologies), using step size of 5 um, at a speed of 2.5 um/sec. Following 1-3 hours of neurophysiological recordings, the probe was retracted at the same step size and speed. The typical insertion depth was 4-9 mm depending on whether the target was the cerebellar cortex or nucleus.

Despite regular cleaning between recording sessions, there was a gradual increase in both the magnitude and phase of the probe’s impedance (Fig. 1A, 1st and 2nd row, linear mixed effects model, main effect of number of days of recording: 22% increase in impedance magnitude per day of recording, t(2110) = 36.2, p = 1.46 e^-223^, 3.06-degree increase in impedance phase per day of recording, t(2110) = 35.1, p = 7.4 e^-213^). Each of the probes had 64 contacts. We characterized a contact as ‘bad’ when it had an impedance magnitude of over 150k Ohms. The number of bad contacts increased with recording sessions (Fig. 1A, third row, linear mixed effects model, main effect of number of days of recording: 4.6- contact increase in bad contacts per day of recording, t(31) = 2.89, p = 0.0069).

**Figure 1.**
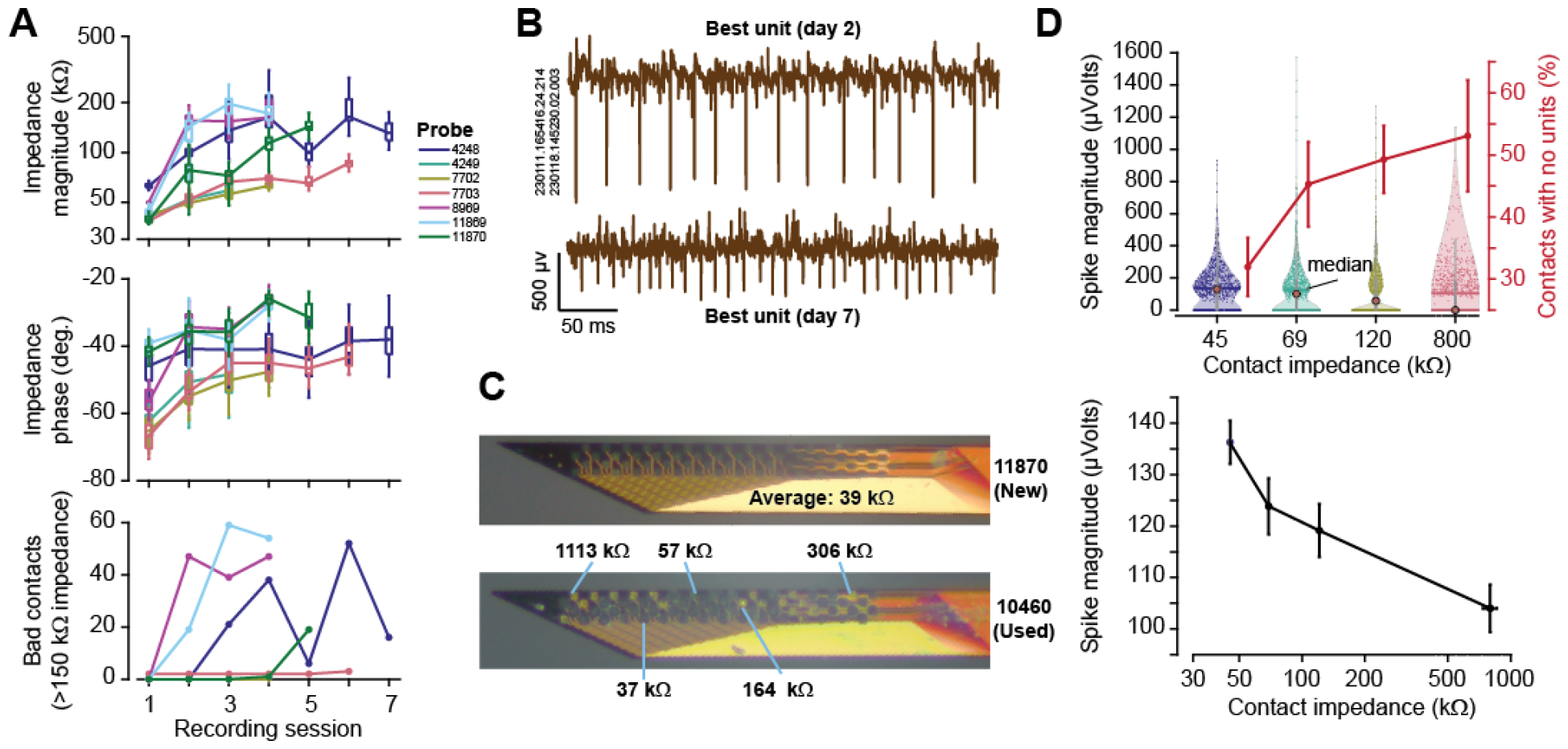
The factory-deposited PEDOT coating degraded across recording sessions, leading to increased impedance and decreased recording quality. **A**. Magnitude, phase, and number of bad contacts (impedance magnitude > 150k Ohms) as a function of recording sessions. **B**. Representative raw data recordings from the best neuron, Day 2 and Day 7. Spikes from the best unit are 2-3 times larger on Day 2 as compared to Day 7. **C**. Microscope images from two probes. Top: new probe 11870, which had an average impedance magnitude of 39k Ohm. The dark blue contacts have good PEDOT coating, characteristic of a new probe. Bottom: used probe 10460, which had contacts of highly variable impedance magnitude. The gold contacts have poor PEDOT coating with a corresponding high magnitude of impedance. Many contacts are on a spectrum between these two extremes due to uneven wear. **D**. Top: the spike magnitude of the signal recorded on a contact vs. the impedance of that contact. The red trace is the percentage of contacts per bin that had no well-isolated units. Error bars indicate variance, and are computed from bootstrapped data (n = 1000 simulated probes). Bottom: spike magnitude as a function of impedance. Error bars are SEM.

Impedance magnitude represents the resistive properties of the contact, where lower magnitude indicates that charge can pass more easily. Impedance phase measures the capacitive properties, where a perfect capacitor has a -90° phase (Macdonald, 1987). The increase in magnitude and phase that we found with repeated use suggested that the charge-carrying capacity of the contacts was decreasing. To understand the cause of these changes, we imaged the probes in their new state, then again after each recording. Fig. 1C illustrates a new and a used probe, along with the impedance magnitude of various contacts. On the new probe, the contacts were dark blue and uniform in color.

This indicated an intact PEDOT coating, matching the measured average impedance of 39k Ohms with a standard deviation of .003k Ohms. The used probe had some contacts that were still well-coated (dark blue, 37k Ohms and 57k Ohms), but also many contacts that were almost bare (164k Ohms, 306k Ohms, 1113k Ohms), exhibiting a light-blue or gold color. This suggested that some of the coating was lost during the recording sessions.

This loss of the coating polymer appeared to affect both the quality and quantity of units that we were able to record. Typically, the spike waveform of the sortable units on the first day of a probe was much larger than the waveform on later recordings. For example, on probe 10460, the best unit on Day 2 was 2-3 times larger than the best unit for the same probe on Day 7 (Fig. 1B). We examined the data over days of recording by plotting spike magnitude as a function of impedance magnitude (Fig. 1D). As impedance increased, spike magnitude decreased, and the percentage of contacts that had no units increased (binned by magnitude of impedance into 4 groups. ANOVA on median spike magnitude per contact for 976 contacts per group, f(3) = 7.46, p = 5.63e^-5^. Significance test for slope of percentage of contacts with no units across 4 groups (top plot, red line), data assigned randomly to each of 4 groups to create null distribution, repeating 400 times. Slope significantly different from null distribution with p < 1e^-5^, z-score = 9.4939. Thus, with repeated use, the probes lost some of their polymer coating, and this coincided with increased impedance and a reduction in signal quality.

### Each recording session deposited tissue on the probe, requiring cleaning that removed the tissue but not the polymer coating

An additional challenge that we faced was that with each recording, there was an accumulation of debris on the probe. Under the microscope the debris appeared as a fibrous tissue containing proteins, dura-gel, and dust from the surrounding air (Fig. 2A, left image). Thus, we needed to clean the probes, but there was a risk that the cleaning method would remove both the tissue and the polymer coating.

**Figure 2.**
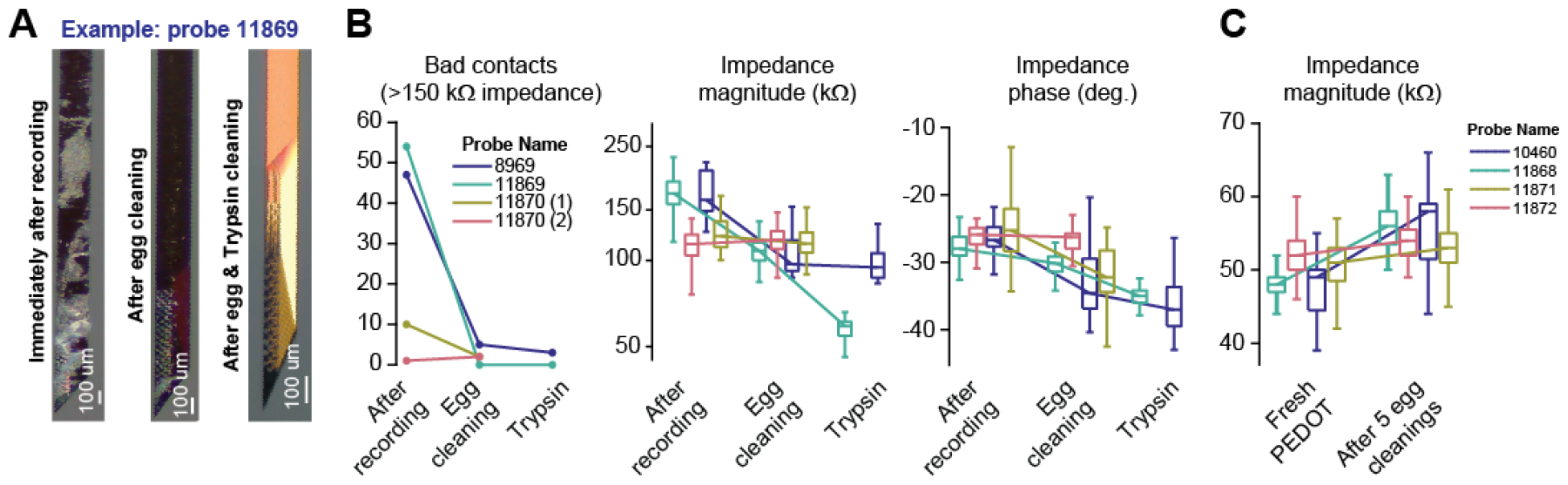
Method for removing tissue that electrophysiological recordings deposited on the probes. The objective was to find a method that removed tissue without also removing the PEDOT coating. **A**. Image of a probe immediately after completion of recording, after inserting into a hardboiled egg, and after soaking in Trypsin. **B**. The effects of egg cleaning and Trypsin on the impedance properties of the various probes. **C**. Effects of egg cleaning on the PEDOT. Error bars are box and whisker plots, indicating the median, upper and lower quartile, and upper and lower extreme.

We tested two conventional methods (enzyme cleaner and Trypsin), as well as an unconventional method of cleaning (hard-boiled egg). Surprisingly, cleaning with a hard-boiled egg followed by Trypsin proved to be the most reliable method. We began by repeatedly inserting the probe into a peeled, hard-boiled egg. This removed most of the tissue that had adhered to the probe (Fig 2A, middle image). Next, we removed the remaining debris, and potentially any debris resulting from the egg cleaning, using an overnight soak in Trypsin (Fig 2A, right image). Trypsin is a solution commonly used to dissociate cells from each other during cell culture but is also used to clean biological matter from probes (Tokiwa et al., 1979; Cui et al., 2001; Cui and Martin, 2003; Alba et al., 2015; Neto et al., 2016, 2018). We quantified the effectiveness of this method by measuring the magnitude and phase of impedance, as well as the number of bad contacts (Fig. 2B), and observed that egg cleaning followed by Trypsin reduced impedance magnitude and phase, as well as the number of bad contacts (linear mixed effects model, main effect of egg cleaning then trypsin: 88% reduction in impedance magnitude, t(638) = -21.1, p = 1.74 e^-75^, 8.2 degree reduction in impedance phase, t(638) = -29.8, p = 6.10 e^-123^, 33.4-contact reduction in bad contacts, t(8) = -2.7, p = 0.027).

A critical question was whether this cleaning method harmed the polymer coating. Trypsin has been documented as a safe cleaning method for silicon probes with PEDOT ((Scott et al., 2012; Alba et al., 2015; Neto et al., 2016, 2018)), but what about the egg cleaning? In order to test if egg cleaning was safe for the PEDOT coating, probes with a fresh PEDOT coating were subjected to 5 egg cleanings. Each egg cleaning consisted of inserting the probe into the egg and moving up and down the shank with a scrubbing motion. The scrubbing motion was repeated for approximately the same amount of time it took to clean a dirty probe, an average of 15 times. This was repeated five times for each probe, representing five days of cleaning between recording sessions. We found little evidence that egg cleaning affected the PEDOT coating (Fig 2C, signed rank test, median increase of 5kOhm over 5 simulated egg cleanings, median absolute deviation of 3kOhm, p = 2.97 e^-37^). The slight increase in impedance may have been due to deposition of egg particles on the probe, which was removed by soaking the probe in Trypsin for an hour. Thus, the combination of hard-boiled egg cleaning and subsequent soaking in Trypsin appeared to be an effective way to clean the probes.

The conventional method of cleaning probes, including tungsten and non-coated silicon probes, is soaking in a solution of enzyme cleaner (Tergazyme, Alconox Inc.). This cleaner can remove proteins and tissue without harming the base material ((Chen et al., 2023)). However, we observed adverse effects using this method, finding that impedance magnitude increased greatly, with many contacts in the mega Ohms range (Figure 3B), an indication of reduced quality of PEDOT coating. For example, on a typical probe there was visible debris and fibers after removal from the brain (Fig. 3A). After an overnight soak in 2% Tergazyme solution with stirring, the probe appeared to still have some debris. This debris was removed with egg cleaning, which revealed that the contacts were gold color, indicating a loss of the PEDOT coating. Therefore, Tergazyme enzyme cleaner removed the PEDOT coating.

**Figure 3.**
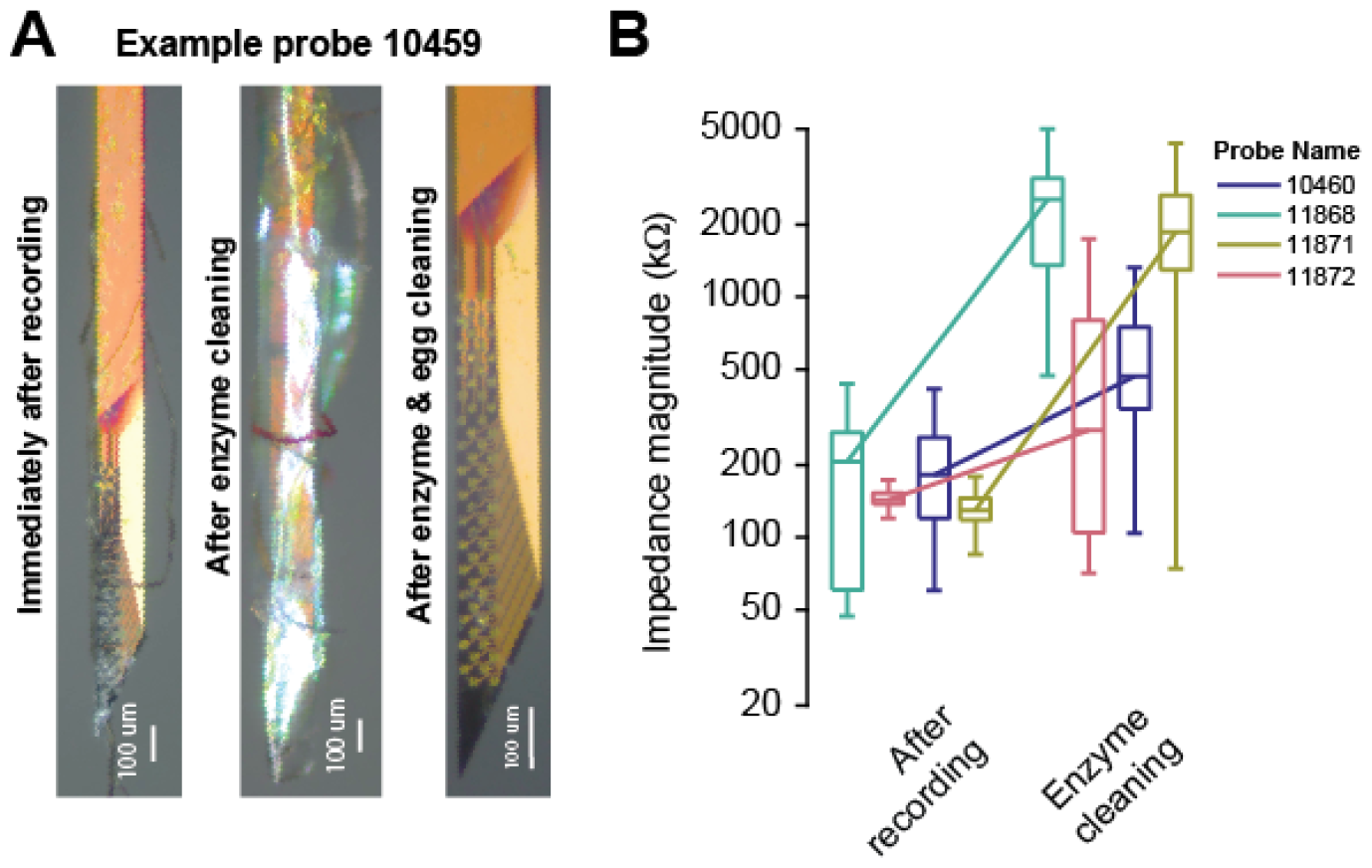
Enzyme cleaning stripped the PEDOT coating. A. Image of a probe immediately after completion of recording, after enzyme cleaning, and after egg cleaning. B. Effect of enzyme cleaning on probe impedance. Error bars are box and whisker plots, indicating the median, upper and lower quartile, and upper and lower extreme.

In summary, enzyme cleaner was an inappropriate method for cleaning these probes because it stripped the polymer coating, dramatically increasing the probe’s impedance. An effective method was hard-boiled egg cleaning followed by Trypsin.

### Recoating via electrodeposition

Despite regular cleaning, insertion into the brain led to gradual loss of the factory-deposited PEDOT coating, resulting in reduced signal quality (Fig 1A). In order to restore the probes, the PEDOT coating needed to be re-deposited. We approached this with electrodeposition, creating a mixture of PEDOT and PSS and using the NanoZ Electroplate mode to plate each contact. However, prior to recoating, we found it essential to remove all PEDOT coating. We did this using Tergazyme enzyme cleaner, and then deposited new PEDOT onto the cleaned gold contacts. This method proved to rejuvenate the probes to like-new condition.

Fig. 4A shows the probe following enzyme stripping, with clean gold contacts (left image). After the probe was recoated with fresh PEDOT, the contacts took on a dark blue-black color, indicating an even coating. The quality of recoating was tested with impedance measurements. Fig. 4B shows the impedances of four probes across the rejuvenation process. These probes started with a median impedance of about 150-200k Ohms, with large variance in impedance across the probes, indicating uneven wear of the PEDOT coating. We next stripped the PEDOT coating with Tergazyme, leaving the contacts clean of any debris and mostly clear of the original factory PEDOT. This greatly increased probe impedance (linear mixed effects model, main effect of enzyme stripping: 483% increase in impedance magnitude, t(510) = 24.0, p = 9.44 e^-86^). The probes were then recoated, resulting in a median impedance at approximately 50k Ohms, which is as good or better than the impedance measurements tested in the new probes. The phase component was restored as well, indicating that the recoated probe had similar resistive and capacitive properties as a new probe.

**Figure 4.**
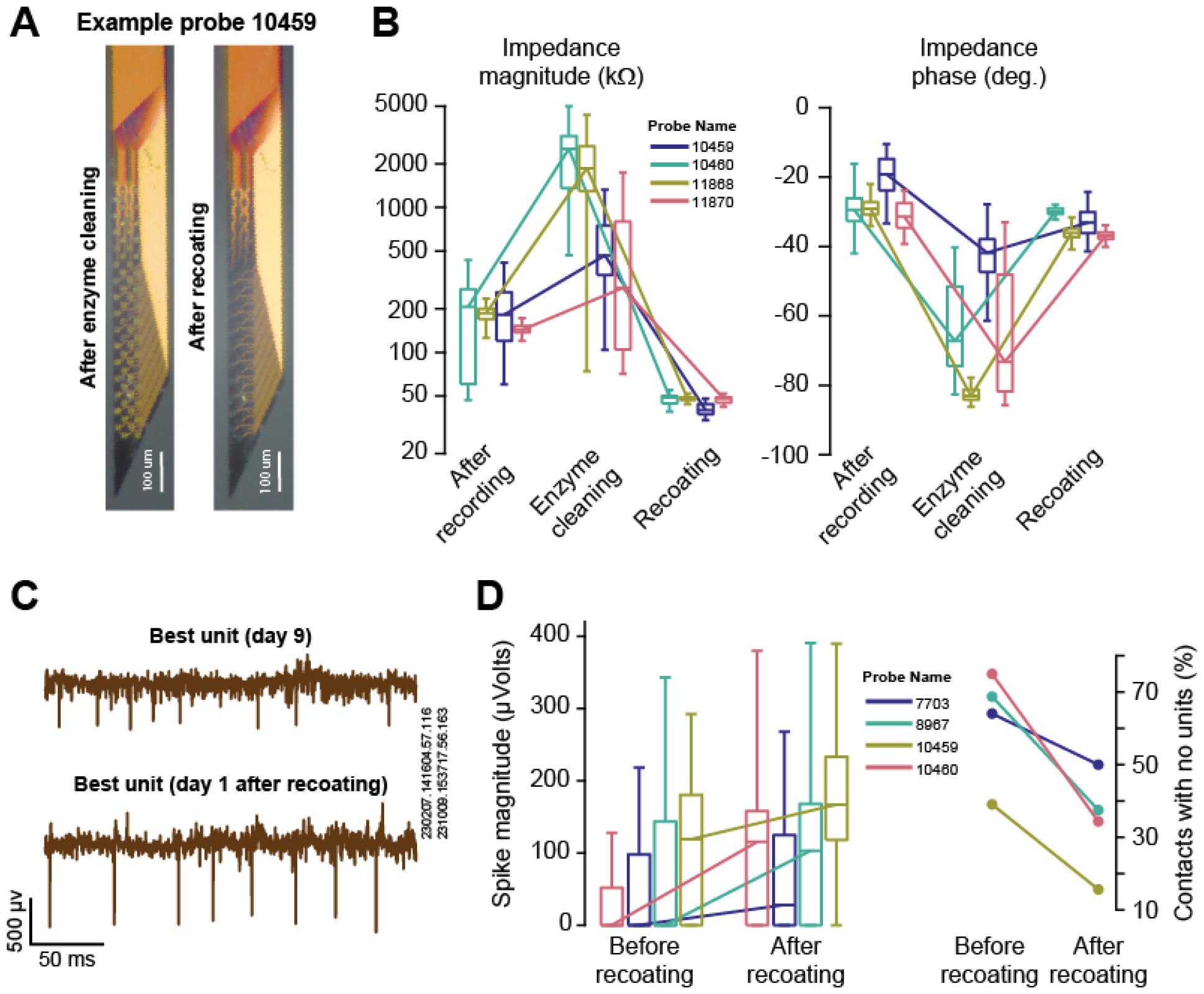
Recoating rejuvenated the probes. **A**. Image of a probe after enzyme cleaning, which stripped the PEDOT coating, and after recoating. **B**. Impedance properties of the probes following recording, enzyme cleaning, and recoating. **C**. Representative raw data from the best unit that was recorded on Day 9, and the best unit that was recorded after the probe was recoated. **D**. Spike magnitude and number of contacts with no units before and after recoating. Error bars are box and whisker plots, indicating the median, upper and lower quartile, and upper and lower extreme.

Although impedance is an indicator of the PEDOT coating quality, we sought to test the rejuvenation method via signal quality of electrophysiological recording. Despite confounding differences such as location of recording in the brain and other factors that affect quality of recording, we found that recoating indeed improved signal quality. Fig. 4C demonstrates data for a single probe on a recording day immediately before and after the recoating procedure. The best unit from the recording day before recoating, when the probe had been used for 9 recording sessions, had spikes that were 2-3 times smaller than the best unit from the first day after recoating. In four probes that we examined before and after recoating, there was a 57.4 uV increase in average spike magnitude of isolated units with recoating (Fig 4D left panel, linear mixed effects model, main effect of recoating: t(510) = 6.12, p = 1.88 e^-9^). In addition, the number of contacts with no isolated units decreased by 27% (Fig 4D, right panel, ANOVA f(1) = 6.65, p = .0419).

In summary, recoating of the conductive polymer returned impedance magnitudes to around 50k Ohms and significantly improved signal quality.

### Methods that did not yield positive results

For cleaning, we first tried soaking the probe in Trypsin-EDTA overnight. This did not remove the bulk of debris from the probe or greatly improve the impedance on the debris-obscured contacts. We also tried changing the order of cleaning to first trypsin, and then hard-boiled egg cleaning. This method was less effective than egg followed by Trypsin, potentially due to the mechanical agitation from the egg loosening all of the debris to a state where any remaining debris could be removed chemically. We also tried cleaning the probe mechanically, with similar technique to the hard-boiled egg method, in a cup of either 1:1 ratio cured dura-gel (Cambridge Neurotech) or prepared 1.5%, 2%, 3%, or 4% laboratory-grade Agar (Innovating Science, Amazon). The dura-gel was sticky but not strong enough, and instead deposited itself onto the probe in small chunks. The agar was not sticky enough, despite being almost too hard to penetrate safely with the probe.

We also tried other methods for recoating the probes. Originally, we attempted plating over the existing PEDOT coating in order to thicken it, only cleaning the probe beforehand with egg or trypsin, or both. However, those methods resulted in uneven coating, with most contacts still in the high-impedance range. This suggests that the existing PEDOT either prevents new PEDOT from adhering to the gold contact, or the existing PEDOT coating is dirty or damaged in a way that prevents appropriate electroplating conditions at the site.

## Discussion

To better understand why the signal quality of some silicon probes degrades with use, we employed Cambridge probes to record from the marmoset brain. On the one hand, insertion into the brain caused a loss of the probe’s PEDOT coating, and on the other hand it deposited tissue on the probe, both of which promoted increased impedance and reduced signal quality. Inserting the probe into a hard-boiled egg followed by soaking in Trypsin was effective in removing the tissue. To replace the lost coating, we first stripped the contacts using an enzyme cleaner, then recoated the probes via electroplating. This cleaning, stripping, and recoating procedure rejuvenated the probes, returning their impedance to around 50k Ohms. Critically, recoating dramatically improved the capacity of the used probes to record electrophysiological signals from the brain.

### Impedance is correlated to spike magnitude and the ability to isolate single units

While low impedance is generally a desirable property in recording electrodes, it is not clear that lower impedance corresponds to improved neuronal signal quality. For example, (Neto et al., 2018) compared bare iridium and PEDOT-PSS coated probes in an anesthetized preparation and found no significant differences in acutely recorded spike magnitudes. This was despite significant differences in impedance and noise level in saline. In contrast, (Baião, 2014) compared uncoated gold and PEDOT-coated gold contacts in acute anesthetized recordings and found that the coated probes not only exhibited reduced impedance, they also had better signal quality. (Cui and Martin, 2003) recorded acutely in anesthetized animals and found larger signal amplitudes in the PEDOT coated contacts as compared to the uncoated contacts, which shared the same significant difference in impedance between coating conditions. In contrast, (Scott et al., 2012) did not find a signal quality difference in lower-impedance gold-coated iridium probes, but did observe a noise reduction. Others have found that in chronic recordings, coated probes produced a greater number of well-isolated units, but the relationship to impedance was unclear because those measurements were taken while the probe was in the brain (Ludwig et al., 2006; Venkatraman et al., 2009; Alba et al., 2015). Taken together, previous work had suggested that uncoated electrodes had a higher innate impedance and lower ability to record units than PEDOT-coated electrodes.

A limitation of many of these studies is that they only compared coated and uncoated electrodes, which usually differ in impedance by one or two orders of magnitude. There is no study currently that directly correlates impedance on a spectrum from well-coated electrodes to a degraded or absent coating. Here, we measured impedance and signal quality on each of 64 contacts in 12 probes across 61 acute recording sessions and found that spike magnitude was inversely varied with impedance (Fig 1D). As impedance increased, our ability to isolate single units greatly diminished. Moreover, because recoating reduced impedance and improved signal quality, the loss of the coating and its associated increase in impedance was likely a causal factor in the reduced signal quality.

### Limitations

Although we used the same charge density for recoating each electrode, and generally recoated the probes to an impedance magnitude of about 50k Ohms, we were unable to directly measure coating thickness. Coating thickness is one of the more important characteristics of conductive polymer coating, and is directly correlated with impedance (Cui et al., 2001; Dijk et al., 2020a, 2020b; Niederhoffer et al., 2023). However, because coating thickness measurements are generally destructive, we did not employ them in our experiments.

Our methods and conclusions are only directly applicable to PEDOT-PSS coating on passive gold contacts for acute recordings in the brain. There is a vast literature on the use of other dopants besides PSS, or other coatings entirely for various applications (Cui et al., 2001; Cui and Martin, 2003; Ferguson et al., 2009; Ludwig et al., 2011; Scott et al., 2012; Baião, 2014; Alba et al., 2015; Carli et al., 2019; Wang et al., 2021; Niederhoffer et al., 2023). However, PEDOT-PSS seems to be an effective and popular conductive polymer coating (Cui and Martin, 2003; Venkatraman et al., 2009; Park et al., 2013; Koutsouras et al., 2017; Neto et al., 2018; Pranti et al., 2018; Boehler et al., 2019; Dijk et al., 2020a, 2020b; Jones et al., 2020).

Our conclusions regarding signal quality were based on acute recordings, which necessarily varied in trajectory and individual experimenter electrophysiology technique among other factors that may influence recording quality or wear on the probe. Despite these confounding factors, within-probe comparisons suggested that use promoted loss of the coating, and recoating improved signal quality.

Perhaps our most surprising result was the observation that inserting the probe into a hard-boiled egg was an effective way to remove some of the tissues that are deposited on the probes. We conjecture that the efficacy is due to the springy but sticky nature of the protein matrix in the egg white, and the way in which it gently but firmly glides along the probe shank. This method does have a drawback in that it is mechanical, not chemical, and thus care must be taken to not break the probe.

## Methods

All data were acquired using 64-contact large animal silicon probes (M1 checkerboard and M2 linear Cambridge Neurotech). Neurophysiological data were collected from three marmosets (*Callithrix Jacchus*, 2 male and 1 female, 350–370 g, between 6 and 8 yrs old, subjects Mirza, Ramon, and Charlie), using methods described earlier (Sedaghat-Nejad et al., 2019). The marmosets were born and raised in a colony that Prof. Xiaoqin Wang has maintained at the Johns Hopkins School of Medicine since 1996. The procedures on the marmosets were approved by the Johns Hopkins University Animal Care and Use Committee in compliance with the guidelines of the United States National Institutes of Health.

### Impedance measurements

At the onset of each electrophysiological recording session, we measured the impedance of each contact by inserting the probe in sterile saline at 1k Hz via an Intan RHD2000-series Data Acquisition (DAQ) System using the OpenEphys (Siegle et al., 2017) Impedance Measurement utility. The electrode was connected to a 64-channel head stage amplifier and digitizer (Intan Technologies). Additional impedance measurements were acquired using a NanoZ impedance device (White Matter LLC.) at a range of frequencies in 0.01M phosphate buffered saline solution (PBS, Millipore Sigma) with a Ag/AgCl (3M KCl) reference electrode (BASi Research Products). We used a 3D-printed holder to maintain a consistent distance between the probe and the reference electrode. If the test signal was clipped at any point during the measurement, the measurement was repeated. Each measurement was an average of 40 sine wave cycles at various frequencies.

### Electrophysiological recordings

Neurophysiological data was recorded acutely in awake, behaving marmosets as they performed a targeted saccade task (Sedaghat-Nejad et al., 2019, 2022). The recordings were from lobules VI and VII of the cerebellar cortex, or from the fastigial nucleus of the cerebellum. Sessions lasted between 1 and 3 hours of recording on any given day, not including penetration and retraction time which varied from 1 to 3 hours total. Thus, the total experiment time was 5-6 hours. Endpoint depth varied between 4 and 9 mm from the skull surface. The probes were inserted through a craniotomy that was covered with a layer of Dura-Gel (Cambridge Neurotech) which protected the exposed dura and supported the probe during penetration. We used a Narishige SM-11 stereotaxic micromanipulator to align and advance into the brain until we reached the cerebellum, at which point we switched to a piezoelectric, high precision microdrive (0.5 μm resolution) with an integrated absolute encoder (M3-LA-3.4-15 Linear smart stage, New Scale Technologies) to advance the electrode to the final recording position.

We connected each probe to a 64-channel head stage amplifier and digitizer (Intan Technologies), and then connected the head stage to the same Intan DAQ that was used for the impedance measurement. Data were sampled at 30 kHz. We used OpenEphys for interfacing with the RHD2000 system and recording of signals. The neurophysiological data were sorted either via P-sort (Sedaghat-Nejad et al., 2021) or via Kilosort 2.0 (Pachitariu et al., 2016) and then curated using the open-source toolbox Phy.

### Cleaning method

During acute electrophysiology, we inserted the probe into the brain, recorded from neurons, and then removed it. This led to deposition of tissue on the probes. To remove this tissue, we tried three methods: inserting the probe into a hard-boiled egg, washing it in a Trypsin solution, and washing it with an enzyme cleaner. For egg cleaning, the probe was mounted onto a micromanipulator and repeatedly inserted into a peeled hard-boiled egg. This process generally consisted of 5-20 cycles of insertion- removal, using a gentle ‘scrubbing’ up-down motion with the manipulator. This process was done carefully to reduce bending of the shank. 70% alcohol was sprayed on the shank between insertions to aid in removal of debris. After each insertion the probe was moved slightly to a new insertion point. Cleanliness was checked between insertions using a microscope, and the process concluded when the probe was visibly free of debris.

For Trypsin cleaning, the probe shank was submerged in Trypsin-EDTA 0.25% solution (Millipore Sigma) overnight. In some cases, the solution was stirred during submersion, but that did not make an appreciable difference as compared to non-stirred submersion.

For enzyme cleaning, the probe shank was submerged in freshly made 2% Tergazyme (Alconox Inc.) dissolved in deionized (DI) water. The solution was stirred continuously over a 12 hour period. If there was visible debris left on the probe, the probe was egg cleaned before taking any measurements. This procedure was repeated once or twice, or until the magnitude of impedance on all contacts was over 300k Ohm and the contacts appeared gold-colored (without PEDOT coating) under the microscope.

### Recoating method

The probes were recoated using a degassed solution of 10 mM EDOT (3,4-Ethylenedioxythiophene, Millipore Sigma) with 32 uM PSS (Poly(sodium 4-styrenesulfonate), Millipore Sigma) in DI water. The NanoZ provided a current source, and utilizing the NanoZ software’s DC electroplating mode, we recoated each contact with a current density of about 3mA/cm^2^ for 30 seconds. The deposition process was repeated on each contact until the impedance dropped to approximately 50k Ohms, although a majority of the contacts only needed one cycle of the process.

### Data Analysis: spike waveform

The magnitude of a neuron’s spike waveform was determined by obtaining an average unit waveform using Phy’s built-in function _*get*_*mean_waveforms*, which averages 100 random spike samples per unit over a roughly 3 ms window. We then computed the peak-to-peak voltage in microvolts from the maximum and minimum values of the average waveform. The value was taken from the contact where the spike waveform was largest.

The best unit for a given day was determined as the largest average waveform size out of all units except complex spikes. Complex spike units were excluded because they are unique to the molecular and purkinje layers of the cerebellar cortex and have a characteristically large peak-to-peak voltage when recording somatically that could bias the data towards having a higher value when recording from specific depths or trajectories. During a given day of data collection, neurophysiology was logged in ∼30 minute durations to reduce file size, so the best unit was taken only from units isolated in the final recording of the session, when the cells were most stable.

For a given day of acute neurophysiology, units were sorted for each ∼30 minute recording. The average unit size was calculated for units found on each recording. Then, the spike magnitude for each contact was calculated as the median size of units found on the given contact for the whole day. If no units were found on a given contact for any recording all day, the magnitude value for that contact is zero.

## Acknowledgements

The work was supported by grants from the NIH (R01-EB028156, R37-NS128416).

